# Interplay between SpaO variants shapes the architecture of the *Salmonella* type III secretion sorting platform

**DOI:** 10.1101/2025.05.15.654217

**Authors:** José Eduardo Soto, Tingting Wang, Jorge E. Galán, Maria Lara-Tejero

**Affiliations:** Department of Microbial Pathogenesis, Yale University School of Medicine, New Haven CT06536

**Keywords:** type III secretion system, *Salmonella enterica*, bacterial pathogenesis, protein secretion

## Abstract

*Salmonella enterica* utilizes a virulence-associated type III secretion system (T3SS) to inject bacterial effectors directly into host cells. Central to this machinery is the sorting platform (SP), a cytosolic assembly whose core scaffolding protein, SpaO, is produced in two isoforms: a full-length (SpaO^L^) and a shorter variant (SpaO^short^) comprising the C-terminal 101 residues of SpaO^L^. Although SpaO^short^ is evolutionarily conserved across type III secretion systems, its precise function has remained elusive. Here, we combined a sensitive, real-time translocation assay with site-directed photo-crosslinking to elucidate the role of SpaO^short^ in *Salmonella* SPI-1 T3SS. We found that while SpaO^short^ is not absolutely required for effector secretion, its absence significantly dampens T3SS-mediated protein delivery. Further biochemical and structural probing revealed that SpaO^short^ is a structural component of the sorting platform, arranged as a homodimer associated to SpaO^L^ via an N-terminal “docking motif.” This interaction occurs while SpaO^L^ is associated with other SP components, supporting a model in which SpaO^short^ is integrated into the SP pods alongside SpaO^L^, OrgA, and OrgB. Collectively, these findings show that SpaO^short^, while not strictly essential, functions as a critical structural component of the sorting platform, providing new insights into how *Salmonella* and related bacteria assemble and maintain these specialized protein-injection systems.

## Introduction

*Salmonella enterica* remains a leading cause of acute gastroenteritis worldwide, posing a significant public health burden particularly in low- and middle-income countries [1]. Its virulence is largely driven by the type <u>III s</u>ecretion <u>s</u>ystem (T3SS), a complex molecular device that delivers effector proteins into target eukaryotic cells [2, 3]. The T3SS or injectisome forms a translocation conduit enabling effector proteins to move from the bacterial cytoplasm into the host cell. *Salmonella* pathogenesis requires the sequential action of two distinct T3SS encoded within the *<u>S</u>almonella* <u>p</u>athogenicity islands 1 (SPI-1) [4, 5] and 2 (SPI-2) [6]. The SPI-1 T3SS mediates early host cell invasion and inflammation, whereas the SPI-2 T3SS supports intracellular survival and replication.

From an evolutionary standpoint, virulence-associated T3SS are believed to have emerged through exaptation of an ancestral flagellar system, wherein motility-dedicated components were repurposed for protein injection [7]. Accordingly, the virulence T3SS shares many structural and mechanistic features with the bacterial flagellar system [8]. This evolutionary adaptation has been independently acquired by many other important pathogens such as *Shigella* spp., *Yersinia* spp., and *Pseudomonas* spp., and confers a competitive advantage by allowing these organisms to interact intimately with eukaryotic host cells.

The T3SS architecture comprises two key structural modules: an envelope-embedded needle complex (NC) and a large cytosolic assembly known as the sorting platform (SP). The NC is a supramolecular structure built of stacked membrane rings anchoring a hollow needle-like projection that extends into the extracellular space [9]. Within the NC, the export apparatus [10], a conical protein assembly partially embedded in the inner membrane, helps form a continuous channel for client protein transport [11, 12]. Attached to the cytosolic side of the NC is the SP [13, 14]. While the NC provides a conduit for effector passage through the bacterial envelope, the SP functions as a hub that orchestrates intracellular processes required for protein injection. Notably, the SP selects, sorts, and initiates client proteins destined for the T3SS pathway, ensuring efficient and timely delivery of effectors into host cells [13].

Recent *in situ* cryo-electron tomography studies in *Shigella* [15] and *Salmonella* [16] have revealed that the SP adopts an intricate, cage-like structure composed of five distinct proteins, each with its own stoichiometry. Together, these proteins form six “pods” arranged radially around a central hub. Each pod connects to the NC via the adaptor protein OrgA, while the SP’s core protein, SpaO, binds OrgA through its N-terminal domain, forming the pod’s main body. At the base of each pod, an OrgB dimer attaches to the SpaO C-terminus, and extends inward to create a cradle for an InvC hexamer, which serves as the ATPase responsible for chaperone release from cognate secreted proteins [17]. This chamber-like enclosure is thought to create a privileged compartment for client protein sorting and unfolding [14]. Although the flagellar system also contains a homologous cytosolic structure known as the C-ring, its architecture is markedly different from that of the SP [18]. Whereas the SP forms a partially open cage-like structure, the flagellar C-ring is a closed structure, likely reflecting its role in torque transmission.

SpaO is the primary constituent of the SP in *Salmonella*. This 303-amino-acid protein acts as a central hub for protein-protein interactions within the SP [19, 20], promoting the formation of the pods. Structurally, SpaO consists of an N-terminal region comprising the first third of the protein, which lacks experimental structural data but is predicted by AlphaFold to form a globular domain, followed by two <u>s</u>urface <u>p</u>resentation of <u>a</u>ntigens domains (SPOA1 and SPOA2). These domains adopt a domain-swapped arrangement critical for numerous protein-protein interactions [21].

A unique and conserved feature of the *spaO* gene, and its homologs in other T3SS-containing bacteria, is that it encodes two distinct products: the full-length SpaO (SpaO^L^) and a shorter isoform (SpaO^short^) comprising the last 101 amino acids. The latter is generated from an internal translation initiation site located at the terminal third of the gene [20]. The presence of a similar short isoform in *Shigellla* [22], *Yersinia* [23], *Xathomonas* [24], and in the *Salmonella* SPI-2 T3SS [25], strongly suggests that SpaO^short^ plays an important role in T3SS function. Unlike SpaO^L^, SpaO^short^ contains only the SPOA2 domain [21], suggesting a function distinct from that of the full-length isoform.

Previous work demonstrated that the SpaO^short^ variant forms tightly intertwined homodimers that bind SpaO^L^ in a 2:1 stoichiometric ratio (two SpaO^short^ per one SpaO^L^) [21, 26]. However, the structural details of this complex have remained elusive. Importantly, it has remained unclear whether SpaO^short^ is a structural component of the sorting platform. Likewise, conflicting data from *in vitro* functional assays [20, 22, 23, 25] have complicated efforts to pinpoint the specific contribution of SpaO^short^ to injectisome assembly and function.

To clarify the elusive function of SpaO^short^, we investigated its functional and structural relationship with SpaO^L^. Using a highly sensitive split NanoLuc-based assay, we quantitatively show that the absence of SpaO^short^ impairs, but does not abolish, T3SS functionality, suggesting an ancillary yet critical role for SpaO^short^ in T3SS activity. Through extensive *in vivo* crosslinking, we uncovered previously unknown interfaces between the two SpaO isoforms, identifying a key structural motif at the SpaO^L^ N-terminus essential for recruiting SpaO^short^. Further *in vivo* protein-interaction studies reveal that SpaO^L^ possesses multiple, non-mutually exclusive binding sites for its partners, findings that have major implications for current structural models of the SP. Overall, this work illuminates how SpaO^short^, an elusive SP component, integrates into both the architecture and function of the T3SS.

### Results

### SpaO^short^ is required for efficient T3SS-mediated protein translocation

To investigate the functional role of SpaO^short^ in T3SS activity, we first sought a method that would not only offer a broader dynamic range and higher sensitivity than standard *in vitro* secretion assays but would also closely mimic the physiological context in which *Salmonella* infects host cells. We therefore adopted a split-nanoLuciferase assay [27], which allows real-time measurement of effector translocation into cultured cells [28, 29]. To implement this assay, we engineered a stable 293 T cell line expressing the large fragment of NanoLuc luciferase, (LgBiT) that lacks a single β-strand critical for its function (here after referred to as LgBiT-293 T). Concurrently, we generated a construct in which the missing β-strand fragment (NP) was fused to the C-terminus of the first 161 aa of the SPI-1 effector SptP (SptP_161_-NP), which includes the secretion signal and the chaperone-binding domain, and co-expressed this construct in *Salmonella*. Functional reconstitution of NanoLuc occurs only if SptP-NP is translocated into host cells via the T3SS, where the NP fragment can complement LgBiT and restore luciferase activity (Fig. 1a). To improve reconstitution kinetics of the split luciferase, we appended the coiled-coil dimer-forming peptides N7 and N8, whose affinities lie in the low nanomolar range [30], to the NP and LgBiT moieties, respectively. The resulting SptP-NP-N7 fusion was expressed from an arabinose-inducible plasmid either in wild-type *S.* Typhimurium, the T3SS-deficient strain Δ*spaO* (negative control), or the Δ*spaO^short^* strain. In the Δ*spaO^short^* strain the internal start codon (GTG_203) for *spaO^short^* is mutated to GCG in the *Salmonella* chromosome, thereby specifically preventing translation of the C-terminal SpaO^short^ isoform [20].

**Figure 1.**
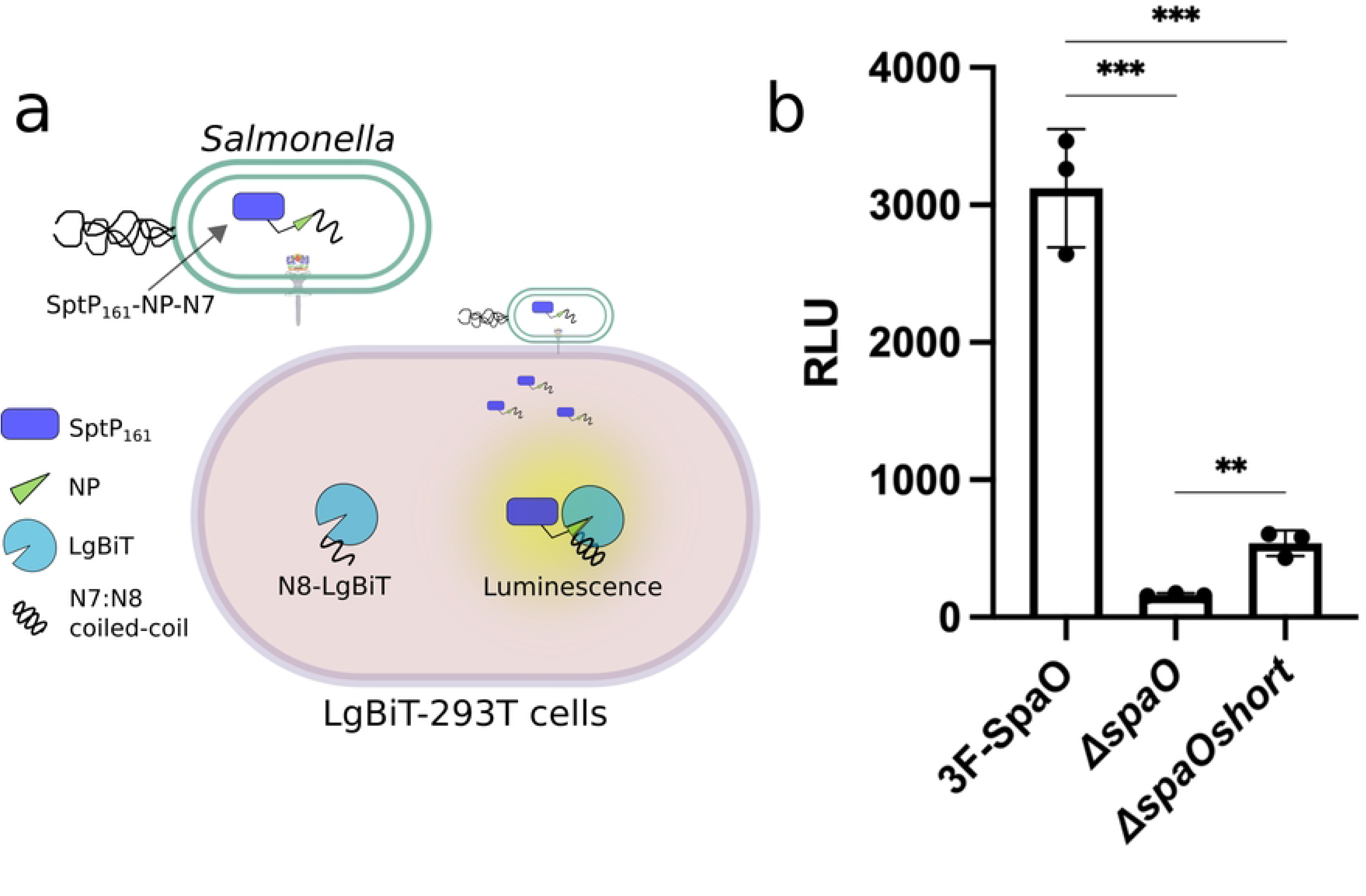
SpaO^short^ is required for fully functional T3SS activity. (a) Schematic diagram of the split NanoLuc-based translocation assay. *S*. Typhimurium was engineered to produce the first 161 residues of the SptP effector fused at its C-terminus to the NanoLuc peptide NP and the high-affinity peptide N7. Host cells stably producing the LgBiT moiety fused to the peptide N8 were infected, and T3SS-dependant translocation of the SptP-Np-N7 effector was detected as a luminescent signal, following reconstitution of the luciferase reporter. (b) Luminescent signal resulting from the translocation of the SptP-NP-N7 effector into LgBiT-293 T cells upon infection with wild-type *S*. Typhimurium or the otherwise isogenic Δ*spaO*, or *ΔspaO^short^* strains. Data expressed as relative light units (RLU) are presented as mean ± standard deviation. Statistical significance was evaluated using a paired Student’s t-test; \*\*\**p* ≤ 0.001; \*\**p* ≤ 0.01.

LgBiT-293 T cells were infected with these strains at multiplicity of infection (MOI) of 1:100 for 1 h allowing T3SS-mediated translocation of SptP-NP-N7. Gentamicin was then added to eliminate extracellular bacteria, followed by washing and cell lysis for luminescence measurements (Fig. 1b). When infected with the wild-type strain, LgBiT-293 T cells produced a robust luciferase signal. By contrast, the luciferase signal was drastically reduced in cells infected with the T3SS-deficient Δ*spaO* strain, confirming the specificity and dynamic range of the assay. Notably, infection with the Δ*spaO^short^*strain yielded a ∼5-fold reduction in signal relative to wild type yet still exceeded the negative control signal. These results indicate that although the absence of SpaO^short^ impairs T3SS function, it does not abolish translocation altogether. Collectively, these findings demonstrate that SpaO^short^, while not essential, plays a pivotal role in ensuring efficient T3SS-mediated effector delivery.

### *In vivo* crosslinking defines the interface architecture of the SpaO^L^-SpaO^short^ complex within the sorting platform

The observed phenotype of the Δ*spaO^short^*mutant prompted us to investigate how SpaO^short^ interacts physically with its binding partner SpaO^L^. Although direct interaction between SpaO^L^ and SpaO^short^ (or their respective homologs) have been extensively documented [20, 22, 23, 25, 26, 31], the precise binding mode remains unclear. SpaO^L^ contains two SPOA domains, SPOA1 (residues 140-218) and SPOA2 (residues 232-303) (Fig. 2a). In contrast, SpaO^short^ comprises solely the SPOA2 domain (Fig. 2a). *in vitro* experiments have shown that the SPOA domains can engage in homotypic (SPOA2–SPOA2) or heterotypic (SPOA2-SPOA1) interactions [21], suggesting the existence of potentially different SpaO^short^/SpaO^L^ interfaces *in vivo*. When and where these interfaces potentially form is unknown. Therefore, to clarify this issue we investigated whether, during sorting platform assembly, SpaO^short^ engages in intermolecular interactions with the SPOA domains of SpaO^L^ (Fig. 2b).

**Figure 2.**
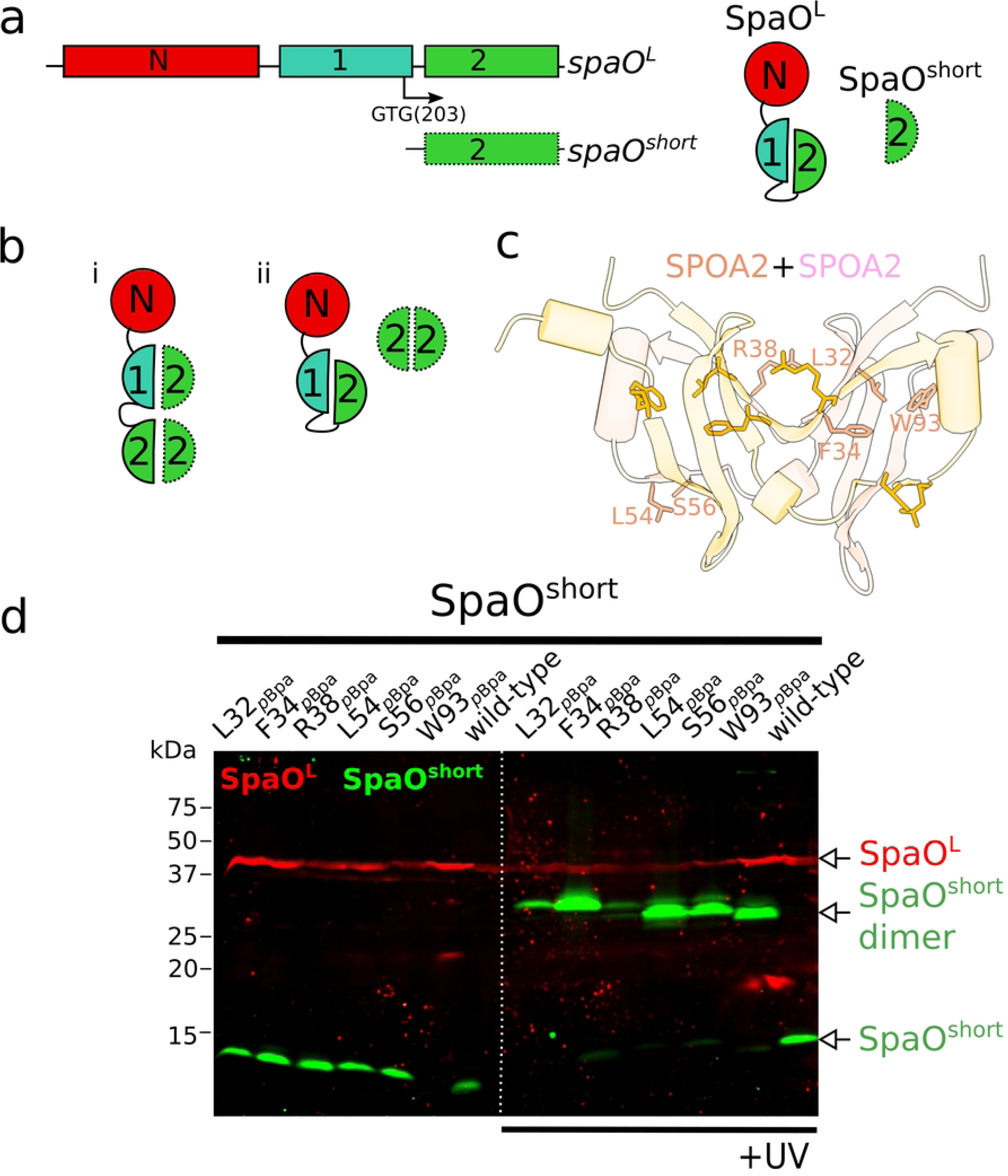
The SPOA2-SPOA2 interface of SpaO^short^ is exclusively committed to dimer formation. (a) Left: domain organization of *spaO^L^* and *spaO^short^*. The internal translation site that leads to the production of SpaO^short^ is marked by an arrow. Right: the SpaO^L^ protein is shown with its N-terminal domain followed by two covalently linked SPOA domains. SpaO^short^ comprises only the SPOA2 domain. (b) Potential interactions models between SpaO^L^ and SpaO^short^. In scenario i, either the SPOA1 or the SPOA2 domain of SpaO^L^ interacts with the SPOA2 domain of SpaO^short^ through the same interface used for SpaO^short^ dimerization. In scenario ii, the SpaO^short^ dimer binds SpaO^L^ using a distinct interaction mode, independent of the intertwined SPOA dimer interface. (c) Solved structure of the SPOA2 domain (PDB 4YX1) displaying the SpaO^short^ residues at the SPOA-SPOA interface that were substituted by *p*Bpa for crosslinking analysis. (d) Immuno-codetection of ^3FLAG^SpaO^L^ (red) and M45-SpaO^short^ (green) from *S*. Typhimurium strains expressing either wild-type proteins as controls, or the indicated *p*Bpa mutant from its native chromosomal locus. Samples were left untreated (left panel) or exposed to UV light (right panel) to induce UV-dependent crosslinking. The detection of the SpaO^short^ crosslinked dimer is indicated.

To this end, we applied a site-specific photo-crosslinking survey informed by the crystal structure of the SPOA2 homodimer [21]. We used the UV-photoactivatable noncanonical amino acid *p*-benzoyl-L-phenylalanine (*p*Bpa), which can be genetically incorporated at any desired position by suppressing a UAG stop codon [32]. Because *p*Bpa has a short reactive distance (∼3.1 Å) [33] that is comparable to typical hydrogen-bond lengths (2.7-3.3 Å) [34], it forms a covalent crosslink only when inserted into actual protein-protein interfaces.

To target SpaO^short^ specifically, we genetically uncoupled *spaO^short^* from *spaO^L^*by replacing the native *spaO* gene with a bicistronic construct coding for 3xFLAG epitope-tagged SpaO^L^ (3xFLAG-SpaO^V203A^) and a downstream, recoded M45 epitope-tagged SpaO^short^ (M45-SpaO^short^), as reported previously [20]. Guided by the SPOA2 homodimer structure (PDB 4YX1), we introduced *p*Bpa at six distinct residues in SpaO^short^ (L32, F34, R38, L54, S56, W93; numbering based on M1 as the first residue of the alternative translation product) located at the dimeric interface (Fig. 2c). UV irradiation of each *p*Bpa-containing strain produced a ∼30 kDa crosslinked species consistent with SpaO^short^ dimer formation (Fig. 2d). However, no SpaO^short^-SpaO^L^ adduct was detected, suggesting that *in vivo* the homotypic SPOA2–SPOA2 interaction is exclusive to SpaO^short^ protomers (scenario ii in Fig. 2b) rather than occurring between SpaO^short^ and any of the SPOA domains of SpaO^L^ (scenario i in Fig. 2b).

Earlier work indicated that, in solution, the SpaO^short^ dimer associates with a single SpaO^L^, resulting in the formation of SpaO^L^-2SpaO^short^ heterotrimer, which can potentially make up the basic building block of each “pod” within the SP [23, 26]. Although it has been previously alternatively suggested that SpaO^L^ engages the SpaO^short^ dimer through either its N-[26] or C-terminal region [20, 25], the precise configuration of this complex has been unclear. To obtain structural insights into the specific arrangement of SpaO^L^ and the SpaO^short^ homodimer complex, we leveraged AlphaFold (AF) [35, 36], which predicted that the first 10 residues of SpaO^L^ anchor the SpaO^short^ dimer through a “docking motif” (Fig. 3a). Specifically, residues 6-8 of SpaO^L^ are predicted to adopt a β-strand conformation when bound to the SpaO^short^ dimer, which aligns with the fifth β-strand of one SpaO^short^ protomer (Fig. 3a, inset). This N-terminal extreme β-strand is not predicted to form when SpaO^L^ is modeled alone, suggesting that SpaO^short^ binding induces SpaO^L^ to adopt this conformation (Fig. 3a, right and Fig. Sup. 1).

**Figure 3.**
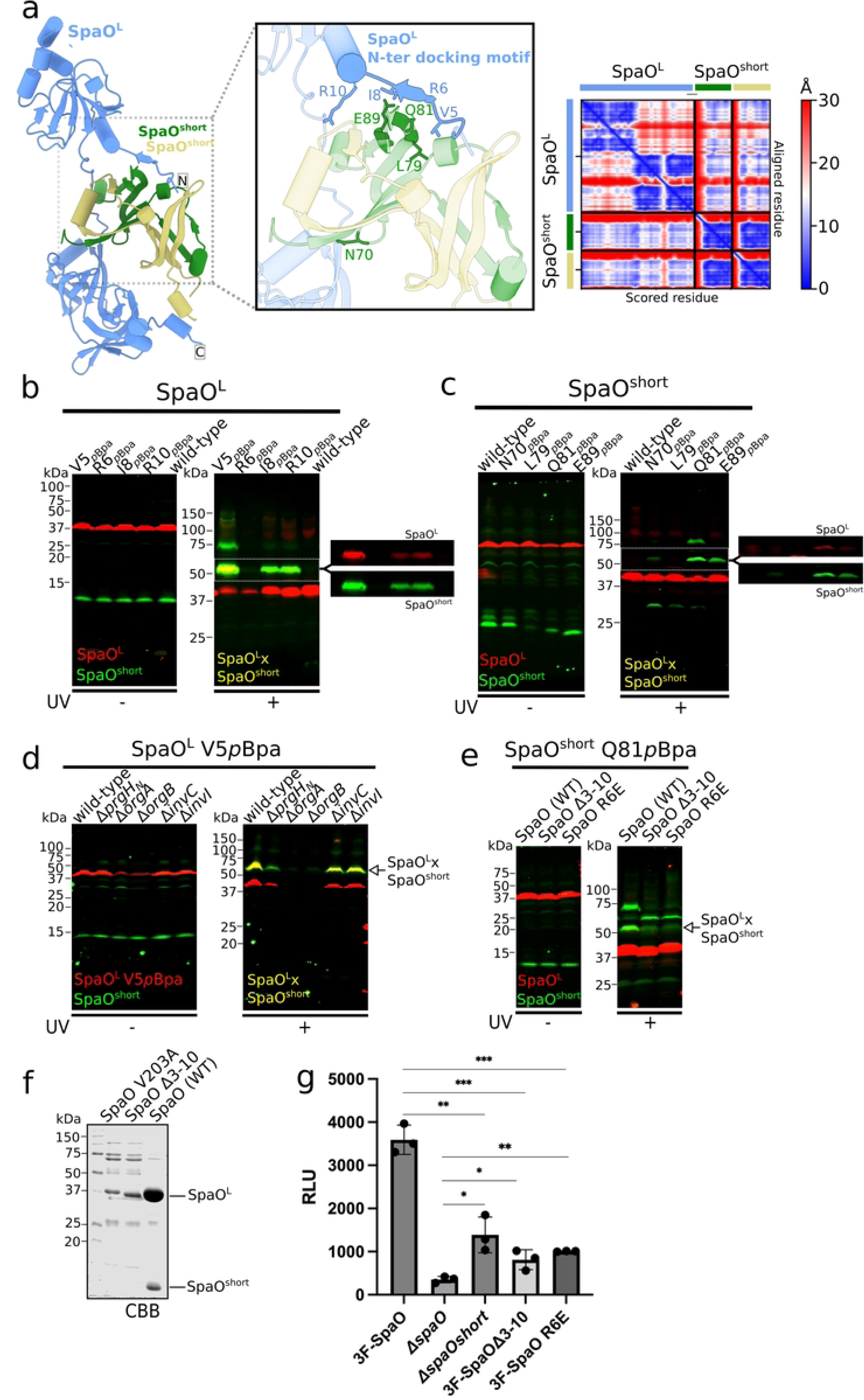
The N-terminal “docking motif” in SpaO^L^ mediates recruitment of SpaO^short^. (a) AlphaFold model of SpaO^L^ (blue) in complex with the SpaO^short^ dimer (green and khaki). (Inset, middle) Close-up view of the predicted docking motif at the SpaO^L^-SpaO^short^ interface. Interfacial residues that were substituted by *p*Bpa for crosslinking analysis are highlighted. The AlphaFold Predicted Alignment Error (PAE) plot on the right indicates the pairwise confidence in residue positioning, ranging from blue (high confidence) to red (low confidence). (b and c) Immuno-codetection of ^3FLAG^SpaO^L^ (red) and M45-SpaO^short^ (green) in *S*. Typhimurium strains expressing either wild-type proteins as controls, or the indicated *p*Bpa-containing SpaO^L^ (b) or SpaO^short^ (c) from their native chromosomal loci. Samples were either left untreated (left) or exposed to UV light (right) to induce UV-dependent crosslinking. The detection of the SpaO^L^-SpaO^short^ crosslinked complex is indicated. (d) Whole-cell lysates of *S*. Typhimurium wild-type or the indicated isogenic mutant strain chromosomally expressing ^3FLAG^SpaO^L^V5pBpa and M45-SpaO^short^, exposed to UV light or left untreated, were analyzed by western blot. The detection of the SpaO^L^-SpaO^short^ crosslinked complex is indicated. (e) Whole-cell lysates of *S*. Typhimurium strains chromosomally expressing M45-SpaO^short^ (green) carrying *p*Bpa at position Q81 (green) and ^3FLAG^SpaO^L^ (red) either wild-type or the indicated mutant derivative. Samples were left untreated (left) or exposed to UV light (right) to induce UV-dependent crosslinking. The detection of the SpaO^L^-SpaO^short^ crosslinked complex is indicated. (f) N-terminal His-tagged SpaO or its mutant derivatives SpaOΔ3-10, or SpaOR6E, expressed in *E. coli*, purified by Ni-NTA resin and visualized by Coomassie brilliant blue staining. His-SpaO^L^ and untagged SpaO^short^ are indicated. (g) Luminescent signal resulting from the translocation of the SptP-NP-N7 effector into LgBiT-293 T cells upon infection with wild-type *S*. Typhimurium chromosomally expressing ^3FLAG^SpaO or the otherwise isogenic Δ*spaO*, Δ*spaO^short^*, ^3FLAG^SpaOΔ3-10, or ^3FLAG^SpaOR6E mutant strains. Data expressed as relative light units (RLU) are presented as mean ± standard deviation. Relevant statistical significance was evaluated using a paired Student’s t-test; \*\*\**p* ≤ 0.001; \*\**p* ≤ 0.01; \**p* ≤ 0.05.

To validate the predicted spatial proximity between the SpaO^L^ N-terminal β-strand (i.e. the “docking motif”) and SpaO^short^, we replaced in the *S.* Typhimurium chromosome four SpaO^L^ residues (V5, R6, I8, and R10) with *p*Bpa in the uncoupled 3xFLAG-SpaO^L^ M45-SpaO^short^ strain. Following UV irradiation of live bacteria, we analyzed crosslinked products by SDS-PAGE and immunoblotting with the anti-FLAG and anti-M45 antibodies. In the negative-control strain (i.e. wild type), which lacks *p*Bpa substitutions, no crosslinks were observed upon UV exposure. However, three out of the four *p*Bpa variants (V5, I8, and R10) produced a robust ∼50 kDa adduct consistent with a SpaO^L^-SpaO^short^ crosslinked complex (Fig. 3b). Fluorescent channel splitting confirmed that the ∼50 kDa species contained both SpaO^L^ and SpaO^short^, demonstrating successful capture of the SpaO^L^-SpaO^short^ complex in *vivo*. Similar crosslinking patterns were detected when the *p*Bpa substitutions were introduced in the native *spaO* locus (i.e., producing SpaO^L^ and SpaO^short^ from a single cistron; Fig. S2) ruling out potential artifacts caused by genetic uncoupling. We noted a ∼75 kDa UV-dependent band in the V5^pBpa^ mutant and in some SpaO^short-pBpa^ substitutions, which we interpret to represent the presence of aberrant migration forms of crosslinked products.

Next, to pinpoint SpaO^short^ residues that interface with SpaO^L^, we carried out “reverse” photo-crosslinking experiments using four surface-exposed SpaO^short^ residues (N70, L79, Q81, E89) that AlphaFold predicts to be in close spatial proximity to SpaO^L^. UV exposure of *p*Bpa-substituted SpaO^short^ variants resulted in the formation of a ∼50 kDa adduct corresponding to SpaO^L^– SpaO^short^, most prominently in Q81^pBpa^ and E89^pBpa^ (Fig. 3c). We also observed a weaker but detectable crosslink in the N70^pBpa^. In the AlphaFold model, Q81 of one SpaO^short^ protomers lies near the docking motif on SpaO^L^ (Fig. 3a, inset), consistent with the strong crosslinking signal. Like what we observed in the V5^pBpa^–SpaO^L^ crosslinks, an additional ∼75 kDa band was detected in the crosslinks of Q81^pBpa^–SpaO^short^, reinforcing the conclusion that these adducts represent the same complex with altered electrophoretic mobility.

With robust crosslink “reporter” sites in hand, we next explored whether formation of the SpaO^L^– SpaO^short^ complex depends on other SP components or a fully assembled T3SS. Introducing the 3×FLAG–SpaO^L-V5pBpa^ allele into strains lacking the ATPase InvC or the stalk protein InvI did not affect the crosslinking patterns (Fig. 3d). Since these mutations do not alter SP assembly but render the injectisome non-functional, these results indicate that the SpaO^L^–SpaO^short^ interaction is independent of an active injectisome. Interestingly, deleting the cytosolic domain of PrgH, which leaves the SP fully cytoplasmic by abrogating its membrane anchor, only mildly reduced SpaO^L^– SpaO^short^ crosslinking relative to the wild-type background (Fig. 3d). In contrast, *orgA* or *orgB* deletions destabilized SpaO^L-V5pBpa^, hindering assessment of the crosslink in those strains. Collectively, these results indicate that SpaO^short^ recruitment by SpaO^L^ occurs prior to the SP’s docking into the needle complex and does not require an intact, functional injectisome.

### The N-terminal docking motif of SpaO^L^ is essential for the recruitment of SpaO^short^ to the sorting platform

To further dissect the functional role of the SpaO^L^ “docking motif,” we examined whether a truncated SpaO^L^ variant lacking residues 3–10 (SpaO^Δ3–10^) could still crosslink with SpaO^short-Q81pBpa^. Deletion of residues 3–10 in *Salmonella* abolished both the ∼50 kDa and ∼75 kDa crosslinked adducts form between SpaO^L^ and SpaO^short-Q81pBpa^ (Fig. 3e). Strikingly, this loss coincided with the appearance of a ∼60 kDa band, which we hypothesize reflects misassembled SpaO^short^ oligomers. Similarly, a point mutation in the docking motif itself, introducing a residue with opposite charge (SpaO^L-R6E^), also disrupted crosslinking with SpaO^short-Q81pBpa^. These data confirm that SpaO^short^ targets the N-terminal β-strand (i.e. the “docking motif”) in SpaO^L^.

We next used *in vitro* pull-down assays to verify the necessity of the N-terminus “docking motif” of SpaO^L^ for SpaO^short^ recruitment. As reported previously [20], heterologous expression of N-terminal His-tagged SpaO^L^ in *E. coli* enables co-purification with SpaO^short^. By contrast, expressing His–SpaO^V203A^ (lacking the SpaO^short^ isoform due to mutation of its internal start codon) markedly reduced the amount of soluble His–SpaO^L^ (Fig. 3f). Notably, His–SpaO^Δ3–10^, which preserves the internal translation start site but lacks residues 3–10, similarly compromised SpaO^L^ solubility and prevented its co-purification with SpaO^short^ (Fig. 3f). These results underscore the pivotal role of the SpaO^L^ N-terminus “docking” motif in SpaO^short^ recruitment.

Because residues 3 to 10 of SpaO^L^ are critical for SpaO^short^ binding, we hypothesized that deleting these residues would phenocopy the functional impact of losing SpaO^short^ entirely. Using the split-nanoLuciferase–based assay, we quantified effector translocation in cells infected by an *S*. Typhimurium strain encoding *spaO^Δ3–10^* on the chromosome. Deletion of residues 3–10 reduced SptP–NP–N7 translocation levels to the same extent observed in the *ΔspaO^short^* strain (Fig. 3 g). In agreement with these results, the SpaO^R6E^ point mutant also exhibited similarly diminished translocation, again mirroring the absence of SpaO^short^. Overall, these findings establish the SpaO^L^ “docking motif” as a crucial determinant for SpaO^short^ recruitment and underscore its essential role in T3SS function.

### Dissection of the modular architecture of the SpaO^L^-SpaO^short^ complex within the sorting platform

SpaO^L^ is the structural scaffold of the *Salmonella* T3SS sorting platform (SP) and acts as a hub for the protein-protein interactions that assemble this megadalton-size complex [16]. Its N-terminal domain binds the symmetry adaptor OrgA [37], whereas the intertwined SPOA1–SPOA2 C-terminal region docks into the OrgB “cradle” [20, 21, 37]. However, the potential structural role of SpaO^short^ within the SP remains unclear. Specifically, it is unknown whether SpaO^short^ is associated with the SpaO^L^ subpopulation engaged in pod formation (i.e. those that contact OrgA and OrgB).

To address this question, we used *in-vivo* photo-crosslinking. The photoactivatable amino acid *p*Bpa was placed at SpaO^L^ residues previously shown to capture SpaO^L^–OrgA (SpaO^T87pBpa^) or SpaO^L^–OrgB (SpaO^N285pBpa^) interactions [37]. These reporters were introduced into a strain that chromosomally expresses OrgA-3×FLAG and M45–SpaO^short^, together with a 2xHA-tagged SpaO^L^ variant carrying *p*Bpa simultaneously at V5 (reporting SpaO^L^–SpaO^short^) and T87 (reporting SpaO^L^–OrgA; Fig. 4a). UV irradiation of this “double *p*Bpa” SpaO^L^ yielded a high-molecular-weight crosslinked adduct that reacted with both anti-FLAG and anti-M45 antibodies, indicating it contained OrgA-3×FLAG and M45–SpaO^short^ (Fig. 4b). Although this band migrated more slowly than expected for a simple trimer, its formation strictly required the simultaneous presence of V5^pBpa^ and T87^pBpa^ in SpaO^L^, showing that a trimeric OrgA–SpaO^L^–SpaO^short^ complex had been captured. Control strains harboring only one *p*Bpa site in SpaO^L^ formed only the corresponding binary complexes (Fig. 4b).

**Figure 4.**
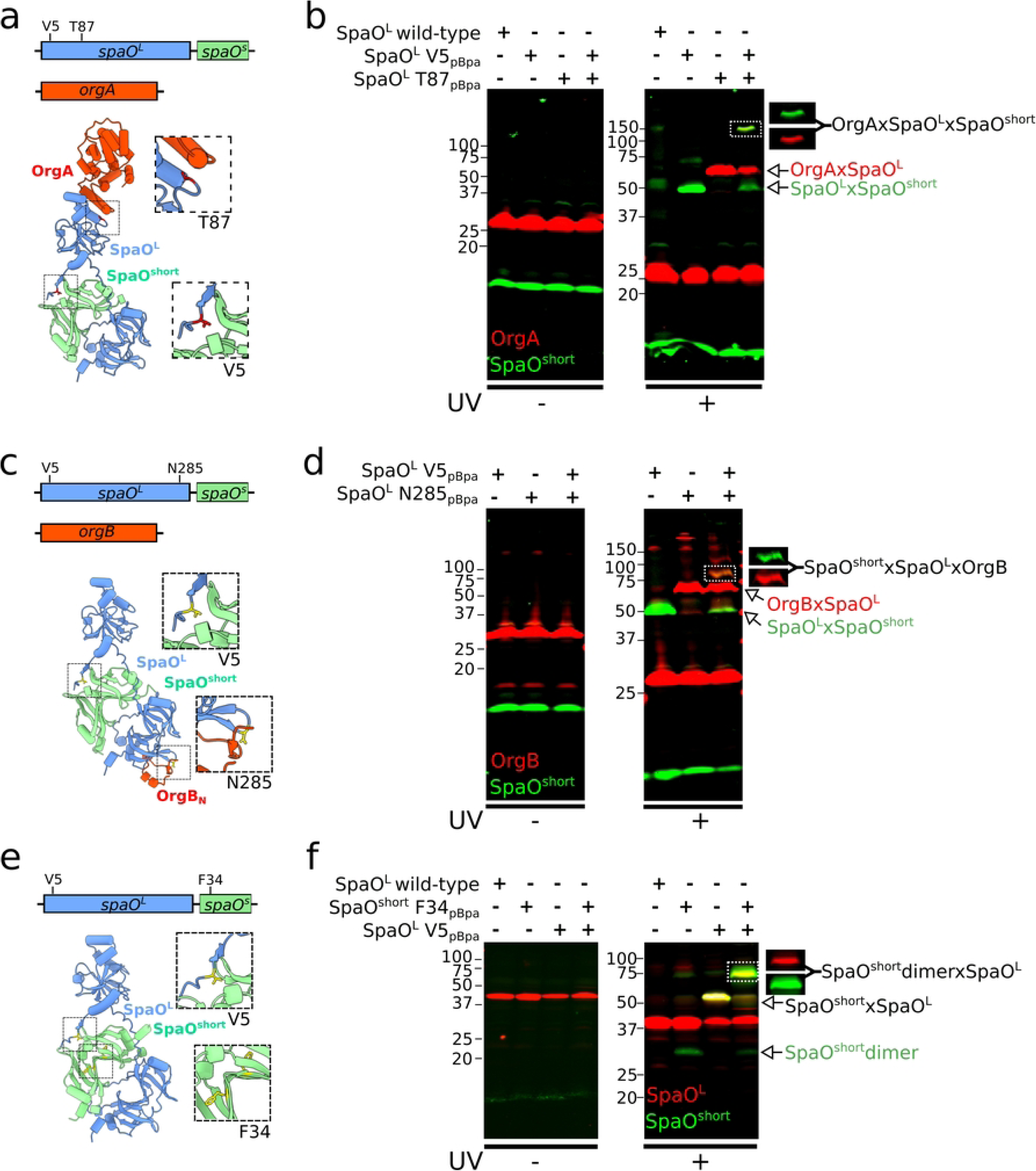
Dual photo-crosslinking reveals the modular architecture of SpaO^L^. (a, c, and e) AlphaFold models of the trimeric complexes (a) OrgA-SpaO^L^-SpaO^short^, (c) OrgB_N_-SpaO^L^-SpaO^short^, and (e) SpaO^L^-SpaO^short^ -SpaO^short^. For the sake of clarity, only the SpaO^L^-binding region of OrgB (OrgB_N_, residues 5-34) is shown in (c). Residues in SpaO^L^ or SpaO^short^ that were substituted by *p*Bpa for photocrosslinking analysis are mapped onto the structure, with close-up views of these regions in the inset. (b, d, and f). Whole-cell lysates from *S*. Typhimurium strains chromosomally expressing (b) M45-SpaO^short^ (green), OrgA-3FLAG (red), and 2xHA-SpaO^L^ V5- and T87-*p*BpA; (d) M45-SpaO^short^ (green), OrgB-3FLAG (red), and 2xHA-SpaO^L^ V5- and N285-*p*BpA; (f) M45-SpaO^short^ (green) F34-*p*BpA (dimer reporter) and 3FLAG-SpaO^L^ (red) V5-*p*Bpa. Samples were left untreated (left) or exposed to UV light (right) to induce UV-dependent crosslinking. The formation of single and double crosslinked adducts are indicated. To show the capture of the trimeric complexes, 700 nm and 800 nm channels were split in the double-crosslinked adducts.

We performed the same analysis with OrgB. A strain encoding 2xHA–SpaO^L^ with *p*Bpa at both V5 (SpaO^short^ binding reporter) and N285 (OrgB binding reporter), was engineered to co-express OrgB-3×FLAG and M45–SpaO^short^ (Fig. 4c). UV exposure produced an ∼80 kDa adduct that was recognized by both, anti-FLAG and anti-M45 antibodies, consistent with a SpaO^L^–OrgB–SpaO^short^ trimer (Fig. 4d).

Together, these experiments demonstrate that a single SpaO^L^ molecule can simultaneously bind SpaO^short^ while simultaneously engaging either OrgA or OrgB. The three interaction surfaces, SpaO^L^ / SpaO^short^, SpaO^L^ / OrgA, and SpaO^L^ / OrgB are therefore spatially distinct and non-overlapping.

To determine whether SpaO^short^ binds SpaO^L^ as a monomer or as a dimer *in vivo,* (i.e. the relative stoichiometry of SpaO^short^ in complex with SpaO^L^), we engineered a strain expressing M45– SpaO^short^ carrying F34*p*Bpa (which reports SpaO^short^ dimerization, Fig. 2c), along with 3xFLAG-SpaO^L^, carrying V5pBpa (which reports SpaO^L^/SpaO^short^ binding, Fig. 4e). Upon UV irradiation, we observed a ∼70 kDa crosslinked adduct containing both M45–SpaO^short^ and 2xHA–SpaO^L^ (Fig. 4f). This result is consistent with SpaO^L^ engaging a preformed SpaO^short^ dimer, supporting a model in which the SpaO^short^ dimer docks onto SpaO^L^ to assemble the functional sorting platform pods.

## Discussion

In this work, we set out to clarify the elusive role of the shorter SpaO isoform (SpaO^short^) in the *Salmonella* SPI-1 type III secretion system. Although SpaO^short^ has long been known to coexist with its full-length counterpart (SpaO^L^) in many T3SS-containing bacteria, its precise function, whether as a structural component of the sorting platform (SP) or as a regulator of its assembly, remained unresolved. By combining a highly sensitive *in vivo* translocation assay, extensive site-directed crosslinking, and structure-guided mutagenesis, we demonstrate that SpaO^short^ is a structural component of the sorting platform, directly engaging SpaO^L^ molecules that are themselves integrated into the SP.

A key finding of our study is that SpaO^short^, while dispensable for T3SS activity in an absolute sense, is nonetheless critical for robust effector translocation into host cells. This result reconciles inconsistencies reported in earlier *in vitro* assays, where the impact of deleting SpaO^short^ in type III protein secretion appeared variable or subtle, by using a more physiologically relevant, real-time approach. Our split nanoLuciferase assay revealed that *Salmonella* strains lacking SpaO^short^ still translocate effectors but at appreciably lower levels than wild type. This suggests a fine-tuning function for SpaO^short^ within the T3SS, likely contributing to the stability and/or efficiency of sorting platform assembly, ultimately enhancing effector translocation.

Building on these functional insights, our photo-crosslinking experiments exposed a previously unknown interface between SpaO^L^ and SpaO^short^, demonstrating that SpaO^short^ is a structural component of the sorting platform rather than a regulatory factor. Contrary to prior hypotheses— such as SpaO^short^ engaging directly with the C-terminal SPOA1–SPOA2 region of SpaO^L^—we found that SpaO^short^ is recruited through an N-terminal “docking motif” in SpaO^L^, which forms a β-strand that stabilizes SpaO^short^ within the sorting platform. Mutations or deletions that disrupt this docking interface prevent SpaO^L^–SpaO^short^ complex formation and impair T3SS-mediated translocation to the same extent as deleting SpaO^short^ outright, underscoring its functional importance. Notably, this arrangement parallels the recently described interaction between the *Salmonella* flagellar proteins FliM and FliN [38], where an N-terminal β-strand extension of FliM binds a FliN dimer [38]. This similarity reinforces the evolutionary link between the T3SS sorting platform and the flagellar C-ring, suggesting that structural principles governing scaffold assembly may be conserved across these two systems.

Our data further demonstrate that SpaO^short^ associates with SpaO^L^ while SpaO^L^ is bound to other key SP components, such as OrgA or OrgB, indicating that these interfaces are not mutually exclusive. This observation directly supports a model in which a single SpaO^L^ molecule can simultaneously interact with multiple partners, reinforcing the concept of SpaO^L^ as a multivalent scaffold. Additionally, by dual-photo-crosslinking both SpaO^L^ and SpaO^short^, we captured a species consistent with a SpaO^L^–2(SpaO^short^) heterotrimer, confirming that SpaO^short^ is incorporated as a homodimer *in vivo*. Together, these findings highlight a sophisticated modular architecture in which SpaO^short^ integrates into the pods of the sorting platform alongside the established OrgA–OrgB– SpaO^L^ framework.

Beyond expanding our basic understanding of T3SS assembly, these insights have several broader implications. First, they help reconcile conflicting phenotypes surrounding SpaO^short^ across different bacterial species by revealing that the shorter isoform fine-tunes effector delivery rather than serving as an essential component. Second, they suggest potential targets for rational therapeutic intervention: compounds that disrupt the SpaO^L^– SpaO^short^ interface could destabilize the entire sorting platform and attenuate virulence without fully inhibiting basic bacterial physiology. Third, our discovery of a conserved docking motif may inform future structural or functional comparisons between T3SSs and the ancestral flagellar system, illuminating how nature repurposes related molecular machinery for distinct roles in motility versus pathogenesis.

In summary, our work establishes SpaO^short^ as an integral structural component of the *Salmonella* sorting platform rather than a regulatory factor. Through a combination of quantitative translocation assays and fine-grained crosslinking, we delineated a new docking motif in SpaO^L^ that recruits a SpaO^short^ homodimer. These findings provide a clearer picture of how the T3SS sorting platform is assembled, offering new perspectives on the molecular organization of this essential virulence system.

## Materials and Methods

### Bacterial strains and plasmids

All strains used in this study are derivatives of *Salmonella enterica* serovar Typhimurium SL1344 [39] and are listed in Table S1. Bacterial strains were routinely grown in lysogenic broth (LB) at 37 °C. When inducing SPI-1 T3SS, *S.* Typhimurium was cultured in LB supplemented with 0.3 M NaCl under low-aeration conditions. Where indicated, expression of the SPI-1 master regulator HilA was induced from an arabinose-inducible plasmid [40]. Selective antibiotics were used at the following concentrations (when required): streptomycin, 100 μg/mL; tetracycline, 10 μg/mL; ampicillin, 100 μg/mL; and chloramphenicol, 10 μg/mL.

Chromosomal modifications (in-frame gene deletions and insertions) were introduced by allelic exchange using the R6K-based suicide vector pSB890 [41] and *E. coli* β-2163 Δ*nic35* as the conjugative donor strain. Plasmids for this work were constructed using Gibson assembly [42] and are listed in Table S1.

### Generation of stable cell lines

A HEK-293 T cell line stably expressing the N8–LgBiT–HA fusion protein was generated by lentiviral transduction followed by puromycin selection, as described previously [43]. Briefly, viruses were produced by co-transfecting HEK-293 T cells with plasmids encoding N8–LgBiT–HA and required packaging factors. After 48 h, the viral supernatant was harvested and used to infect HEK-293 T cells. Transduced cells were then selected in 2 μg/mL puromycin (Gibco) for 5 days, and resistant cells were cloned into 96-well plates to obtain stable single-cell clones.

### Split NanoLuc-based translocation assay

To construct strains for the T3SS-dependent translocation assay, a fragment containing the *sicP* cognate chaperone followed by the first 161 codons of the SPI-1 effector SptP was amplified. The nanoluciferase small fragment (NP) plus the coiled-coil module N7 were fused in frame to the C-terminus of SptP_161_ (SptP–NP–N7). The fusion was cloned into pBAD24A and introduced into the *S. Typhimurium* strains of interest.

HEK-293 T cells stably expressing N8–LgBiT–HA (LgBiT–293 T) were seeded in 24-well plates and grown for 24 h prior to infection. Bacterial cultures, grown under T3SS-inducing conditions (LB + 0.3 M NaCl) with 0.1% arabinose, were added to the LgBiT–293 T cells at a multiplicity of infection of 100 in Hank’s balanced salt solution (Gibco 14025092) containing calcium and magnesium. After a 1 h infection, cells were washed three times with prewarmed PBS and incubated in DMEM (10% bovine calf serum, 50 μg/mL gentamicin) for 1 h to kill extracellular bacteria. Following incubation, cells were washed three additional times with PBS and lysed in H₂O (to avoid lysing intracellular bacteria). Lysates were clarified by centrifugation (14,000 × *g* for 5 min), and the supernatant was transferred to 96-well black-wall plates (Costar). Luciferase activity was measured using the Nano-Glo Luciferase Assay System (Promega) on a Spark multimode microplate reader (Tecan).

### *In vitro* pulldown assay

*E. coli* Lemo21(DE3) strains carrying pSB3775, pSB4539, or pSB2835 were grown overnight and subcultured into 200 mL LB with kanamycin and chloramphenicol at 37 °C until OD_600_ ≈ 0.6. Protein expression was induced with 0.5 mM IPTG, and cultures were incubated for an additional 4 h. Cells were harvested by centrifugation at 6,000 × *g* and resuspended in 5 mL lysis buffer (50 mM NaH₂PO₄, 300 mM NaCl, 10 mM imidazole, 1 mM MgCl₂, 2.5 U/mL DNase, plus cOmplete™ Protease Inhibitor Cocktail [Sigma 11697498001]). Lysis was performed using a One Shot cell disruptor (Constant Systems Ltd., Northants, UK), and cell debris was removed by centrifugation. The cleared lysate was incubated with 100 μL Ni-NTA resin (Qiagen 30210) for 1 h at 4 °C with gentle rocking. Resin was washed four times with 5 mL ice-cold PBS + 20 mM imidazole to remove unbound proteins. Bound proteins were eluted in 200 μL PBS + 200 mM imidazole. Eluted fractions were analyzed by SDS–PAGE and Coomassie Brilliant Blue staining.

### *In vivo* photo-crosslinking

Site-specific incorporation of the photo-crosslinkable amino acid *p*-benzoyl-L-phenylalanine (*p*Bpa, Bachem) was achieved by replacing the chosen codon with a TAG stop codon in the chromosome. This chromosomal strategy avoids artifacts associated with multicopy plasmid expression. The resulting *S. Typhimurium* strains harboring TAG mutations were co-transformed with pSupBpa and pSB3292, enabling TAG suppression by *p*Bpa incorporation and boosting SPI-1 expression, respectively. Overnight cultures of these co-transformants were diluted 1:20 into LB + 0.3 M NaCl containing 50 μg/mL ampicillin, 10 μg/mL chloramphenicol, 1 mM *p*Bpa, and 0.075% arabinose, then grown at 37 °C under low-aeration conditions for 6 h. Cells (1 mL) were transferred to 35-mm tissue-culture dishes (Falcon 353001) and exposed to UV light (λ = 365 nm) from a handheld lamp for 45 min. Control samples were kept in the dark. After crosslinking, cells were pelleted, resuspended in 100 μL SDS–PAGE loading buffer, and 20 μL was run on SDS– PAGE for immunoblot analysis.

### Immunodetection

Protein samples were separated by SDS–PAGE and transferred to nitrocellulose membranes, which were blocked in TBS plus nonfat dry milk. Primary antibodies used were anti-FLAG (Sigma), anti-M45, and anti-HA (BioLegends). Secondary antibodies conjugated to DyLight 800 or DyLight 680 were used to visualize bands on a Li-Cor Odyssey infrared imaging system.

### AlphaFold modeling

An AlphaFold2 [35] structural model of SpaO^L^ in complex with a SpaO^short^ dimer was generated using ColabFold [36]. The model with the lowest predicted alignment error (PAE) was visualized and analyzed in ChimeraX [44]. Inter-residue distances were examined using the SELECT command in ChimeraX.

## Authors contributions

J.E.S. conducted most of the experiments; T.W. developed the luciferase-based translocation assay; M.L.T. and J.E.G provided intellectual input, conceived and directed the project; J.E.S., M.L.T. and J.E.G wrote the manuscript with comments from all the authors.

## Data and materials availability

all data are available in the main text, supplementary materials, and auxiliary files.

## Figure legends

**Fig S1. AlphaFold metrics for the SpaO^L^-2SpaO^short^ complex.** (a) Left, AlphaFold predicted structure of the SpaO^L^ (blue)-SpaO^short^ dimer (green and khaki) complex. Right, the AF per residue confidence score (pLDDT) was mapped onto the structure. The N- and C-terminal ends of SpaO^L^ are indicated. Inset shows zoom-in on the interface between SpaO^L^ docking motif and SpaO^short^. (b) Heatmaps showing the distance in Å between each residue of SpaO^L^ and its nearest residue of SpaO^short^, either in protomer B (upper panel) or C (lower panel). Predicted Aligned Error (PAE) values for the corresponding residue pairs are overlaid. Lower PAE values indicate high confidence relative positioning. Highlighted in red is the identified docking motif in the N-terminal extreme of SpaO^L^.

**Figure S2. Capture of the SpaO^L^-SpaO^short^ crosslinked complex within the native *Salmonella* locus.** Whole cell lysates of *S*. Typhimurium chromosomally expressing either the N-terminally 2HA-tagged wild-type SpaO protein or an isogenic strain expressing the SpaOV5*p*Bpa allele were analyzed by western-blot. Samples were exposed to UV light or left untreated and probed with antibodies against the HA epitope. The detection of the SpaO^L^ -SpaO^short^ crosslinked complex is indicated.

